# A confidence-based reinforcement learning model for perceptual learning

**DOI:** 10.1101/136903

**Authors:** Matthias Guggenmos, Philipp Sterzer

**Affiliations:** Department of Psychiatry and Psychotherapy, Charité Universitätsmedizin Berlin Charitéplatz 1, 10117 Berlin

**Keywords:** Reinforcement learning, perceptual learning, confidence, feedback, exploration

## Abstract

It is well established that learning can occur without external feedback, yet normative reinforcement learning theories have difficulties explaining such instances of learning. Recently, we reported on a confidence-based reinforcement learning model for the model case of perceptual learning (Guggenmos, Wilbertz, Hebart, & Sterzer, 2016), according to which the brain capitalizes on internal monitoring processes when no external feedback is available. In the model, internal confidence prediction errors – the difference between current confidence and expected confidence – serve as teaching signals to guide learning. In the present paper, we explore an extension to this idea. The main idea is that the neural information processing pathways activated for a given sensory stimulus are subject to fluctuations, where some pathway configurations lead to higher confidence than others. Confidence prediction errors strengthen pathway configurations for which fluctuations lead to above-average confidence and weaken those that are associated with below-average confidence. We show through simulation that the model is capable of self-reinforced perceptual learning and can benefit from exploratory network fluctuations. In addition, by simulating different model parameters, we show that the ideal confidence-based learner should (i) exhibit high degrees of network fluctuation in the initial phase of learning, but reduced fluctuations as learning progresses, (ii) have a high learning rate for network updates for immediate performance gains, but a low learning rate for long-term maximum performance, and (iii) be neither too under-nor too overconfident. In sum, we present a model in which confidence prediction errors strengthen favorable network fluctuations and enable learning in the absence of external feedback. The model can be readily applied to data of real-world perceptual tasks in which observers provided both choice and confidence reports.

## 1 Introduction

A well-established form of learning without external feedback is perceptual learning – the ability to improve sensory information processing through training or repeated exposure. Previous models of perceptual learning have successfully used principles from reinforcement learning to explain perceptual learning, but they were limited to cases of external feedback (Kahnt, Grueschow, Speck, & Haynes, 2011; Law & Gold, 2008). Recently, we introduced a novel perceptual learning model, in which learning was guided by internal confidence-based reinforcement signals (Guggenmos, Wilbertz, Hebart, & Sterzer, 2016). Using an associative reinforcement learning rule – a combination of Hebbian learning and confidence prediction errors – the model tuned network weights to improve perception. The model successfully explained perceptual learning behavior in human participants and found dopaminergic source and target regions as neural correlates of confidence-based learning signals, further corroborating a reinforcement learning account of perceptual learning in the absence of external feedback.

However, a simplifying assumption of this previous model was that the variability of confidence reports given constant stimuli was modeled as noise. Here we offer a different treatment of this aspect with the key idea that variability in confidence reports is directly related to network fluctuations in the processing of a stimulus. More specifically, we propose that sensory processing networks employ slightly different pathways configurations for repeated presentations of the same stimulus, where some configurations lead to higher perceptual confidence than others. In the brain, such fluctuations may arise either due to the probabilistic nature of neural information processing or due to an explicit exploratory capacity of neural circuits. Learning in our model is guided by confidence prediction errors that serve to strengthen pathway configurations for which fluctuations led to above-average confidence and to weaken those associated with below-average confidence outcomes. The aim of this work is to introduce the formulation of such a model, to assess its learning behavior and to investigate the influence of key model parameters.

## 2 Model

We consider a classic perceptual learning task, in which observers have to identify whether a noisy Gabor stimulus is oriented counterclockwise (ccw) or clockwise (cw) with respect to a reference axis. The model consists of two sensory input nodes, which represent energy detectors *E*_ccw_ and *E*_cw_ and two decisional output nodes *A*_ccw_ and *A*_cw_. These input and output layers are fully connected with signal weights *w*_ccw,ccw_ and *w*_cw,cw_ connecting input and output nodes with same orientations and noise weights *w*_ccw,cw_ and *w*_cw,ccw_ connecting nodes of opposing orientations. Importantly, the specific processing of a given stimulus is subject to trial-to-trial fluctuations. Fluctuations are implemented by means of a trial-dependent multiplicative (balancing) factor *b*. For instance, the output activities *A*_ccw_ and *A*_cw_ for a counterclockwise stimulus are:

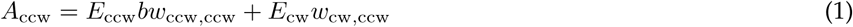

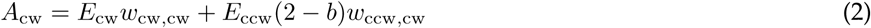

The factor *b* thus shifts the balancing between counterclockwise stimulus energy being processed via the signal pathway (*w*_ccw,ccw_) or the noise pathway (*w*_ccw,cw_).

Confidence *c* is computed as the absolute value of the difference between *A*_ccw_ and *A*_cw_:

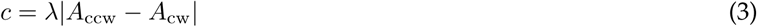

where *λ* is the confidence sensitivity parameter. Learning is guided by a confidence prediction error *δ*, defined as the difference between current confidence *c* and expected confidence 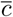:

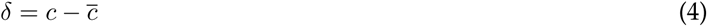

Expected confidence is estimated via a Rescorla-Wagner learning rule:

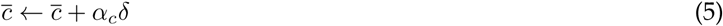

Finally, weights are updated by means of an associative learning reinforcement learning rule:

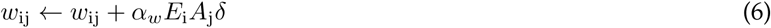

To prevent unlimited growth of the weights, synaptic normalisation is performed:

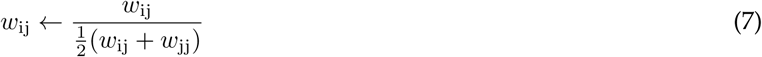

Choices are based on a softmax action selection rule:

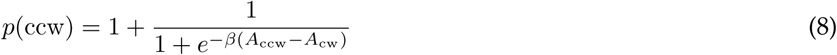

A key feature for empirical application (e.g., model fitting to behavioral data) is that the trial-by-trial balancing factor *b* can be computed from confidence reports by solving Equation (3) for *b*. For counterclockwise stimulus presentation, the analytic derivation for *b* is as follows:

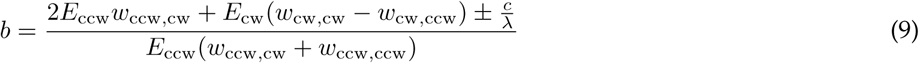

Caused by the absolute value in Equation (3), a single ambiguity remains for b in the term ±*c*/*λ*. As a heuristic, we recommend to choose whichever sign produces a *b* with the smallest deviation from 1, thus slightly biasing the model towards a smaller influence of the balancing component.

## 3 Simulation results

To assess the behavior of the model and its dependence on parameters, we repeatedly confronted the model with 5000 trials of clockwise or counterclockwise stimulus presentations. Each parameter combination was simulated for 10000 runs. Stimuli were a mixture of the designated and the opposing stimulus orientation with a ratio of 3:1. To make the orientation discrimination task challenging, white Gaussian noise was added. The balancing factor *b* was drawn from the following uniform distribution:

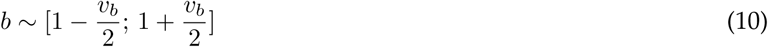

where *υ_b_* determined the magnitude of trial-to-trial weight fluctuations.

For all simulations, if not otherwise specified, the following parameters were chosen: *υ_b_* = 0.5, = 1, *α_w_* = 0.2, *α_c_* = 0.2, *β* = 3. Signal weights were initialized with 1.25, noise weights with 0.75, and initial expected confidence was set to 0.

### 3.1 Successful perceptual learning in the absence of external feedback

Figure 1 shows the the behavior of the model with the above default parameters. Within the first 2000 trials, signal weights have increased and noise weights have decreased to their respective saturation level. As expected from such a tuning of the network, the proportion of correct responses increased as a consequence (from around 0.59 to 0.78). Since confidence was computed from the output activities of the model, confidence and expected confidence likewise showed an increase during learning. Overall, this simulation demonstrated that the model was capable of learning, achieved by an increase of signal weights and a decrease of noise weights.

**Figure 1:**
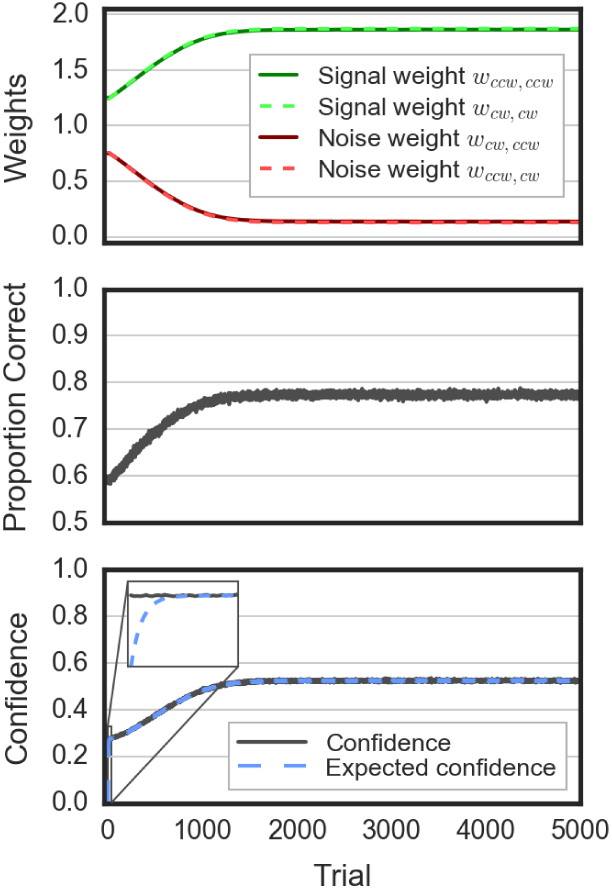
Behavior of the confidence-based perceptual learning model. Learning is achieved by an increase of signal weights and a decrease of noise weights. As a consequence of the weight tuning, performance (proportion correct responses), confidence and expected confidence increase until saturation is reached.

### 3.2 Model parameters affect learning speed and saturation performance

Three parameters directly influenced learning: the magnitude of weight fluctuations (determined by *υ_b_*), the weight-related learning rate *α_w_* and the confidence sensitivity parameter *λ*, which represented the under/over-confidence of the observer. In the following, we discuss the learning behavior of the model in dependence of each of these parameters.

#### 3.2.1 Magnitude of weight fluctuations

The parameter *υ_b_* was the degree to which the network explored different weight configurations. In the initial phase of learning (trials 1-1000), learning was fastest for intermediate values (*υ_b_* ∼ 1) (Figure 2A). Too low values of *υ_b_* led to slower learning as the exploration of weight configurations was negligible and thus had no noticeable positive effect. Too high values of *υ_b_* often induced counterproductive weight configurations, erroneously leading to high confidence for wrong choices. While there was thus a certain sweet spot of *υ_b_* for the initial phase of learning, saturation performance generally decreased with higher *υ_b_*, as the explorativeness became a disadvantage once the network had learned.

**Figure 2:**
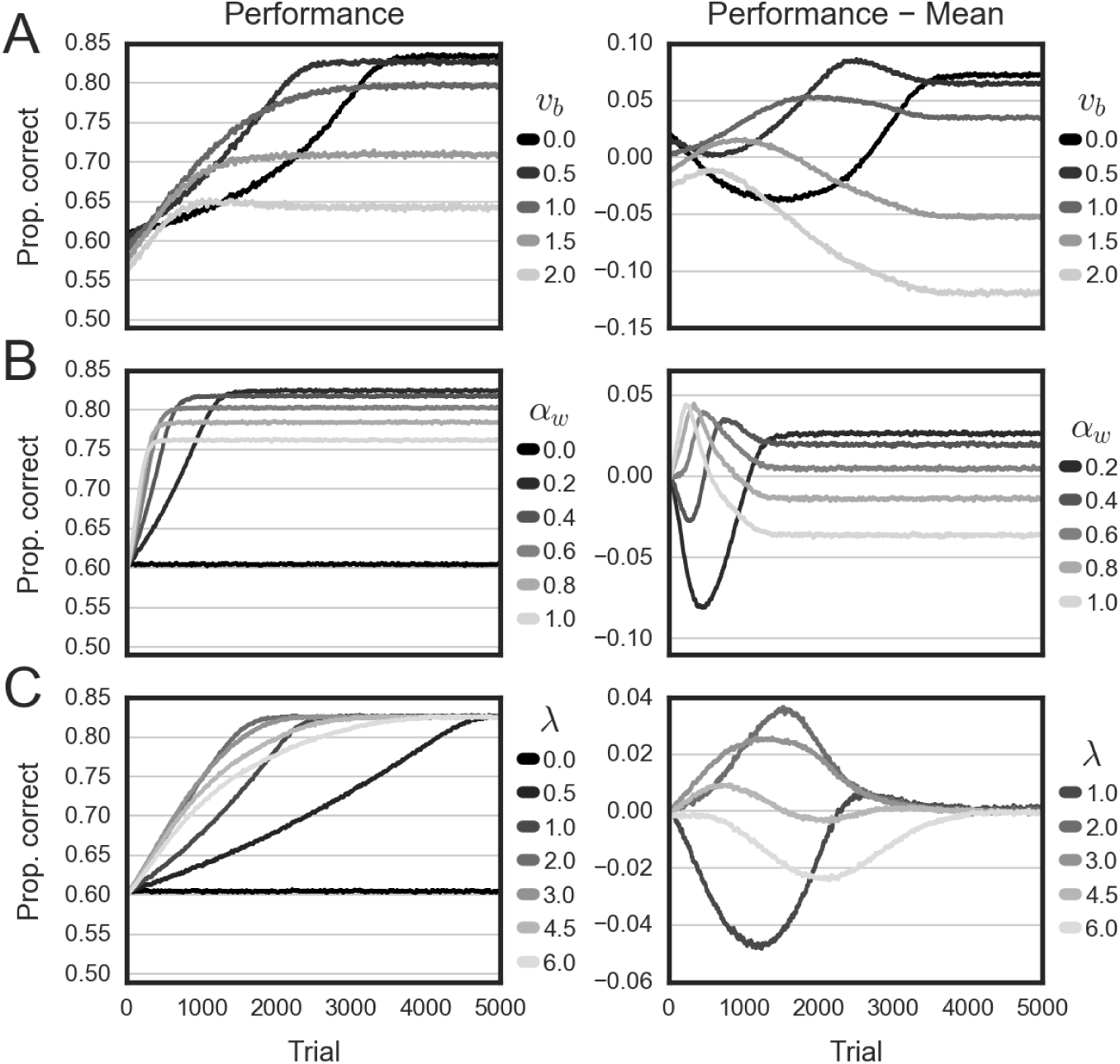
Performance in dependence of model parameters. Left panels show the proportion of correct responses across trials for different parameter settings. In right panels, the mean across parameters was subtracted, for better illustration. **(A)** Magnitude of weight fluctuations *υ_b_*. **(B)** Weight-related learning rate *α_w_*. **(C)** Confidence sensitivity *λ*.

#### 3.2.2 Weight-related learning rate

The parameter *α_w_* determined how much the network learned from confidence prediction errors. In the initial phase of learning, higher learning rates generally led to faster learning (Figure 2B). However, in the long run high learning rates led to suboptimal saturation performance, implying a long-term advantage for slow but steady learners.

#### 3.2.3 Confidence sensitivity

The parameter *λ* determined the degree of confidence for a given activation difference between counterclockwise and clockwise output activities. In the initial phase of learning, fastest learning was achieved for intermediate values (*λ* ∼ 3) (Figure 2C). Too high values of *λ* implied overconfidence and led to overhasty, often disadvantageous, updates of the network. Too low values of *λ* led to tiny confidence-based learning signals and thus reduced learning. Yet, long-term performance levels were almost unaffected by *λ*.

## 4 Conclusions

Here we introduced a novel computational model for perceptual learning in the absence of external feedback, utilizing network fluctuation in combination with confidence prediction errors.

We showed through simulation that the model was capable of learning by increasing signal weights and decreasing noise weights. Importantly, we found that the exploratory network component, implemented as trial-to-trial fluctuations of the weight matrix, was able to boost perceptual learning in the initial phase of learning. However, once the network has reached a stable level, large network fluctuations became a disadvantage and an ideal observer should thus decrease network exploration. This pattern of results mirrors the classic exploration-exploitation dilemma, in which ideal agents have to balance exploration and exploitation depending on their outcomes and the dynamics of the environment.

We further showed that ideal confidence-based l earners s hould n either b e t oo u nderconfident no r to o overconfident for optimal initial learning, while under/overconfidence had little effect on long-term performance levels. By contrast, for the weight-related learning rate, initial gains in performance due to high learning rates came at the cost of reduced long-term performance.

In the future, we plan to apply the model to real-world perceptual learning datasets of human participants in order (i) to assess the model evidence in comparison to prior models, and to (ii) investigate potential individual differences related to the degree of network fluctuations, and how those relate to learning.

## Acknowledgements

This work was supported by the German Research Foundation (Deutsche Forschungsgemeinschaft, DFG, FOR 1617: grants STE1430/6-1 and STE1430/6-2)

